# Parcellation of motor sequence representations in the human neocortex

**DOI:** 10.1101/419754

**Authors:** Atsushi Yokoi, Jörn Diedrichsen

## Abstract

While previous studies have revealed an extended network of cortical regions associated with motor sequence production, the specific role of each of these areas is still elusive. To address this issue, we designed a novel behavioural paradigm that allowed us to experimentally manipulate the structure of motor sequences representations in individual participants. We then conducted fMRI while participants executed 8 trained sequences to examine how this structure is reflected in the associated activity patterns. Both model-based and model-free approaches revealed a clear distinction between primary and non-primary motor cortices in their representational contents, with M1 specifically representing individual finger movements, and premotor and parietal cortices showing a mixture of chunk, sequence and finger transition representations. Using model-free representational parcellation, we could divide these non-primary motor cortices into separate clusters, each with a unique representation along the stimulus-to-action gradient. These results provide new insights into how human neocortex organizes movement sequences.

## Introduction

The highly-sophisticated performance of a pianist often makes us wonder how the brain learns, stores, and produces such complex action sequences. One prevalent idea is that such movements are organized in a hierarchical fashion, where several elementary movements are combined into one single unit, often called “motor chunk” (Lashley, 1951). These motor chunks send ordered descending commands to the generating circuits for each elementary movement with one go, allowing faster and more accurate execution of entire sequence using less cognitive resource (Rosenbaum et al., 1983). These motor chunks could also be further organized into larger chunks or flexibly re-used to create different sequences (Sakai et al., 2003). Such hierarchical structure would greatly reduce computational cost for planning and executing long motor sequences (Ramkumar et al., 2016). An alternative idea is that sequences are *non-hierarchically* represented as a continuous set of transition probabilities between neighbouring movements (Hunt and Aslin, 2001; Reber, 1967; Stadler, 1992; Verwey and Abrahamse, 2012). To date, there is solid behavioural evidence for both organisations. Evidence of whether and how they are represented in the brain is, however, still inconclusive.

Although recent evidence from human functional magnetic resonance imaging (fMRI) studies has suggested that the network of widespread brain regions, including prefrontal cortex (PFC), dorsal/ventral premotor cortex (PMd/v), supplementary motor area (SMA), precuneus, basal ganglia (BG), and areas along intraparietal sulcus (IPS), are *involved* in the production and acquisition of more complex sequences (Grafton et al., 1995; Hikosaka et al., 1999; Honda et al., 1998; Penhune and Steele, 2012; Sadato et al., 1996; Wymbs et al., 2012), the critical question that still remains is *how* sequences are represented in these areas (Hikosaka et al., 1999). Similarly, a number of electrophysiological studies in non-human primates have shown that some neurons show differential firing rates during the same elementary movement, depending on the sequential context such as preceding or following movement (Baldauf et al., 2008; Tanji and Shima, 1994). Other neurons have been found to be active at both initiation and termination of a sequence (Fujii and Graybiel, 2003), or were selective for specific categories of sequences (Shima et al., 2007). However, the use of relatively short and simple sequences in these studies precludes further assessment of hierarchical movement representations.

The aim of this study was therefore to dissociate hierarchical and non-hierarchical sequence representations at the behavioural level, and then investigate their neural correlates. To this end, we used representational fMRI analysis (Diedrichsen and Kriegeskorte, 2017) to investigate how the brain represents a set of well-learned, complex movement sequences (11 finger presses). Rather than analysing the increases or decreases of spatially smoothed activity, representational fMRI analysis makes inferences based on the similarity (or dissimilarity) of multivariate activity patterns across multiple experimental conditions (Ban and Welchman, 2015; Chikazoe et al., 2014; Ejaz et al., 2015; Kriegeskorte et al., 2008; Yokoi et al., 2018). One potential problem in applying this method to study the hierarchical organization of movement sequences is that the specific organisation of motor memories may be different across individuals (Jimenez et al., 2011; Ramkumar et al., 2016), and is often influenced (and hence confounded) by biomechanical constraints (Koch and Hoffmann, 2000). To address this problem, we first established a new behavioural paradigm that allowed us to experimentally manipulate the structure of motor sequence representations.

## Results

### Declarative chunking leads to specific structure of motor memory

Our paradigm was aimed at inducing a stable way of motor chunking by manipulating the way that participants built up their explicit, declarative memory of the sequence. We then sought to detect neuronal correlates of this organisation in the structure of activity pattern in motor and premotor areas as measured with fMRI. During scanning, we required participants to produce sequences with their right hand completely from the memory provided only with a sequence cue (Fig. 1A-C). The training was therefore devoted to enabling participants to remember the 8 sequences (Fig. 1B; for more detail, see Materials & Methods). On day 1, they practiced to produce 8 different chunks of 2 or 3 items (Fig. 1C), and associate these with a specific letter (A-H). On the second day, participants started to learn 8 different sequences as combinations of four of the learned chunks. At the end of this training, participants could reliably recall most of the sequences – the number of incorrect presses they made in each sequence execution had reduced dramatically; the error rate per press was 9±8% (Fig 1D). The inter-press intervals (IPIs) within each chunk quickly reduced on the first day and remained relatively stable for the following days. In contrast, IPIs for the boundaries of two successive chunks were much longer and reduced only slowly over the course of training days (Fig. 1D). Even on the fifth day, the between-chunk IPIs were executed more slowly (439±107ms) than the within-chunk intervals (215±53ms, *t*_12_ =-7.07, *p*=1.31×10^-5^; Fig. 1d). The longer between-chunk IPI is commonly taken as a behavioural indicator that the two sequence elements are stored as separate units.

**Figure 1.**
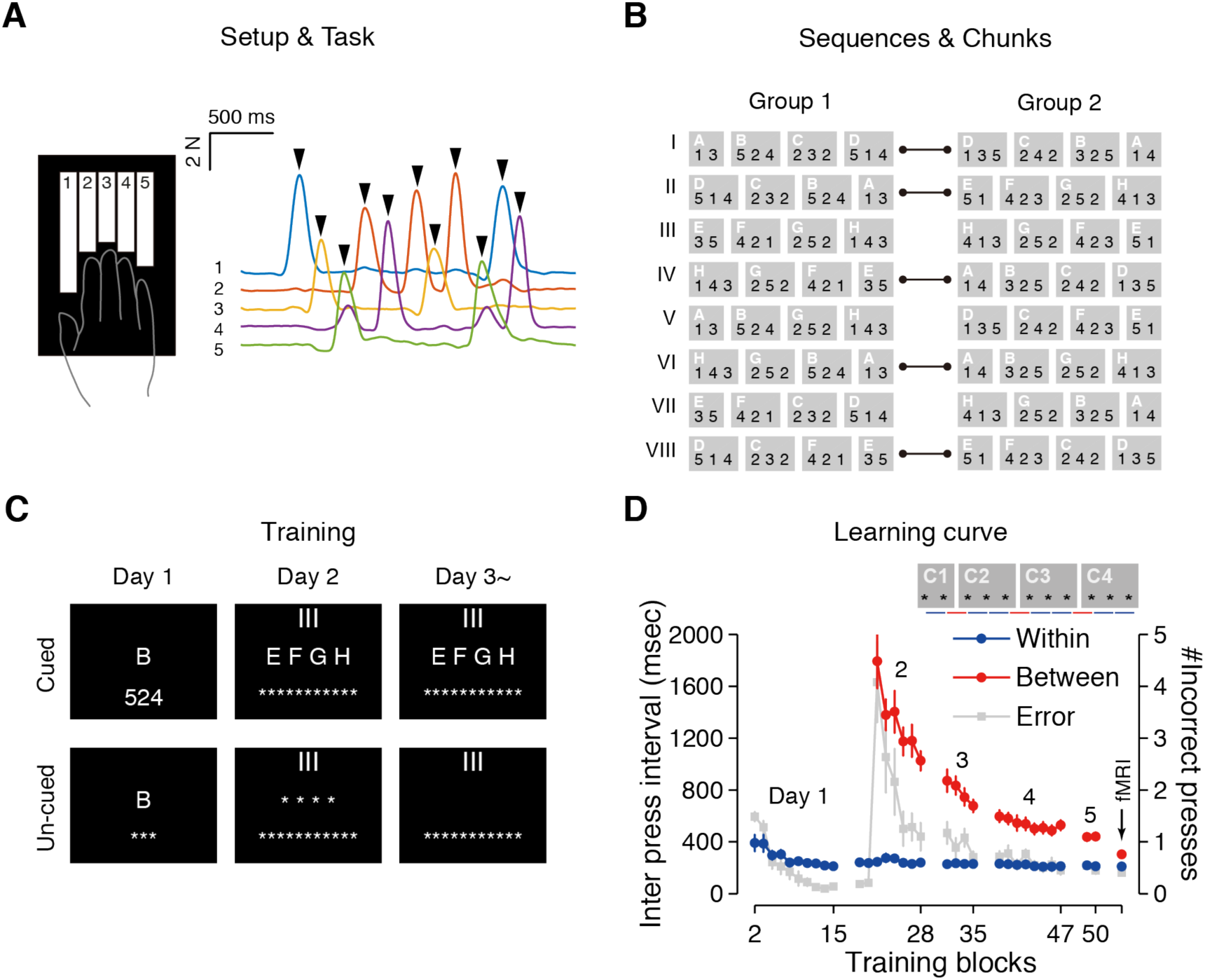
Explicit induction of a structured sequence memory. **(A)** Experimental setup. Participants practiced fast sequence of isometric finger presses on the custom-built keyboard device (left). The traces show example force data (fingers 1-5) from one sequence execution (right). Each arrow-head on peak force indicates a successful finger press. **(B)** Sequences and chunks. The participants were divided into two groups, which practiced partly overlapping sequences with different chunking. Pairs of identical sequences across the two groups are indicated by lines. **(C)** Training consisted of cued trials (upper row) and un-cued trials (lower row). On day 1, participants learned to produce single chunks from memory using a letter (A-H) cue. On the following days, they practiced sequences (indicated by Roman letters I-VIII) as combinations of learned chunks. On Day 2, cued and un-cued trials were alternated. On Day 3-5, cued trials and a set of three un-cued trials were alternated. **(D)** Inter-press intervals over the course of the behavioural training. Within-and between-chunk intervals averaged over the sequence types are displayed in the blue and the red dots, respectively (axis on the left side). Average number of incorrect presses are indicated as the grey squares (axis on the right side). Data points with an arrow was the average performance in the imaging session. Only data from uncued trials are shown. Error-bars indicate the s.e.m.

While this result provided clear evidence for chunking, it remains unclear to what degree the longer between-chunk IPIs are caused by memory retrieval and/or stable motor representation (i.e., planning-ahead of successive movements). To test this, we asked participants to perform a follow-up session after the fMRI scan in which the sequences were instructed not by the sequence cue, but rather directly using all digit cues (e.g., 13524232514, Fig. 2A). This frees participants from the requirement to recall the sequence from memory. Nonetheless, on trained sequences the between-chunk intervals were still longer than the within-chunk intervals (377 ±136ms vs. 275 ±75ms; *t*_14_ = −6.01, *p* = 3.2×10^-5^, Fig. 2E - trained), suggesting that induced chunking affected motor performance over and beyond explicit recall.

**Figure 2.**
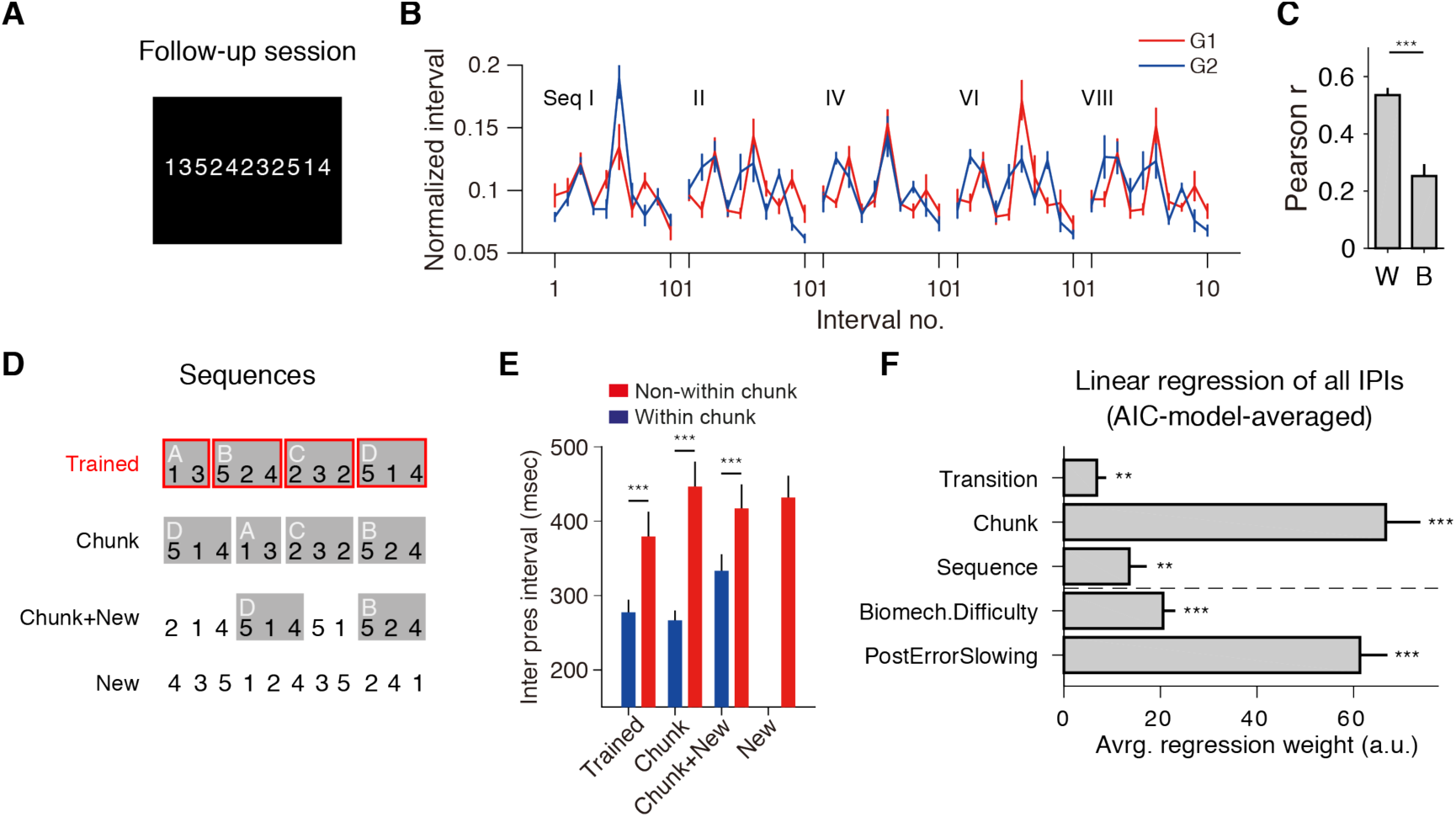
Cognitively induced chunking structure influences subsequent skilled motor performance. **(A)** In the follow-up session conducted after the imaging session, the sequences were directly cued with 11 digits, removing the need for memory recall. **(B)** Average IPIs, normalized to the entire sequence duration, for group G1 and G2. Each group of participants showed group-specific pattern of press intervals, which cannot be attributed to biomechanical difficulty of the finger transitions. **(C)** Correlation (Pearson’s r) of IPI profiles across participants was higher for participants within a group than across the two groups. **(D)** Generalisation test with trained sequences, sequences that consisted of trained chunks in a novel order, sequences that contained trained chunks in random sequences, and completely novel sequences. **(E)** Within-chunk intervals were faster than between-chunk intervals for all sequence categories. (**F**) Group-averaged regression weights for the 3 experimental effects, and 2 nuisance effects. Weight estimates were averaged across all possible regression models using Bayesian model-averaging (see methods). Dashed horizontal line separates between effects of interest and no interest. Error-bars indicate s.e.m. across the subjects. Significance of statistical test were shown by asterisks (*: p<0.05, **: p<0.005, ***: p<0.0001).

Importantly, our data also shows that the observed effect is not driven by differences in the biomechanical difficulty of finger transitions. The sequence design (Fig. 1B) was such that 5 out of the 8 sequences were the same across the two experimental groups, with the difference that they learned them using a different chunking structure. The normalized press intervals for those sequences showed a pattern that clearly reflected the original instruction (Fig. 2B). The inter-subject correlation of these patterns was significantly higher for within-group than between-group comparison (Fig. 2C, *t*_14_ =5.35, *p*=1.0×10^-4^).

If participants acquired a motor representation of the chunks, they should also be able to use this knowledge when producing the chunk in a novel context (Sakai et al., 2003). To test this, the participants additionally executed 3 sets of novel sequences (Fig. 2D). These new sequences either consisted of trained chunks in new combinations and orders (“Chunk”), of 2 trained chunks embedded in an otherwise random sequence (“Chunk+New”), or completely untrained sequences with no relation to learned chunks (“New”). As expected, IPIs in the “New” sequences were executed considerably slower than for IPIs in trained sequences. For the other two sequence types, the intervals that lay within a learned chunk were performed significantly faster than novel intervals (*t*_14_ <-6.01, *p*<3.2×10^-5^; Fig. 2D). Overall, these results suggest that originally declaratively (i.e., cognitively) imposed chunk structure left a reliable imprint in the motor behaviour, which generalized to novel contexts.

Interestingly, we also observed that the between-chunk IPIs of the trained sequences were faster than the between-chunk IPIs of the “Chunk” sequences (*t*_14_ =-3.80, *p*=0.002). This advantage may have two reasons. First, participants may have acquired a higher-order sequence representation that encoded the transitions between chunks. Alternatively, it may be due to a form of non-hierarchical, association learning (Hunt and Aslin, 2001; Reber, 1967; Stadler, 1992; Verwey and Abrahamse, 2012) at the level of the individual elements: finger transitions that had been encountered in practice frequently would become associated and therefore performed faster. To disentangle these two explanations, we modelled all IPIs for the follow-up session for each individual, using three effects of interest: The frequency of this 2-finger transition in training to capture associative learning, whether the interval was within or between chunk to determine the influence of chunking, and whether the chunk transition was already learned within a trained sequence to capture higher-order sequence representations. The full model also contained two effects of no interest: biomechanical effect of each 2-finger transition, and post-error slowing (Botvinick et al., 2001) (see methods). Using Bayesian model-averaging, we determined the influence of each component in the context of the other ones.

The result presented in Fig. 2F indicated both association of individual finger presses and a hierarchical representation of chunk transitions coexisted. Again, we observed very robust within-chunk effect (*t*_14_ =8.19, *p*=1.04×10^-6^, group-*t* test), as well as significant effect of executing known chunk transitions (*t*_14_ =3.51, *p*=0.003), providing a robust behavioural evidence that our participants developed hierarchical representation of the learned finger sequences. Simultaneously, we also found a significant frequency-dependent effect of 2-finger transitions (*t*_14_ =3.90, *p*=0.0016). Thus, our results provided clear evidence for a co-existence of hierarchical and non-hierarchal (associative) representation of movement sequences. With this behavioural evidence, we next assessed where and how these different representations are implemented in different brain regions.

### Cortical regions with robust sequence “encoding”

In the fMRI session, the participants received a brief visual cue for the sequence type and then executed the sequence twice (Fig. 3A). The activation associated with each sequence was estimated for each voxel by averaging the task-evoked BOLD activity over the two executions. We then applied representational fMRI analysis to study the cortical sequence representation. Using a searchlight approach (Fig. 3B) we first determined whether the activation patterns had any information about the executed sequences. For this, we computed a cross-validated estimate of the Mahalanobis distance (crossnobis distance estimator, Diedrichsen et al., 2016; Walther et al., 2016) between any possible pair of the sequences. Systematically positive crossnobis estimates indicate reliable differences between the activity pattern for different sequences. Given that we averaged the activity across two executions and given that all sequences consisted of the same finger presses arranged in a different order (Fig. 1B), any difference in activity patterns must reflect some dependency of the activity on the sequential context.

**Figure 3.**
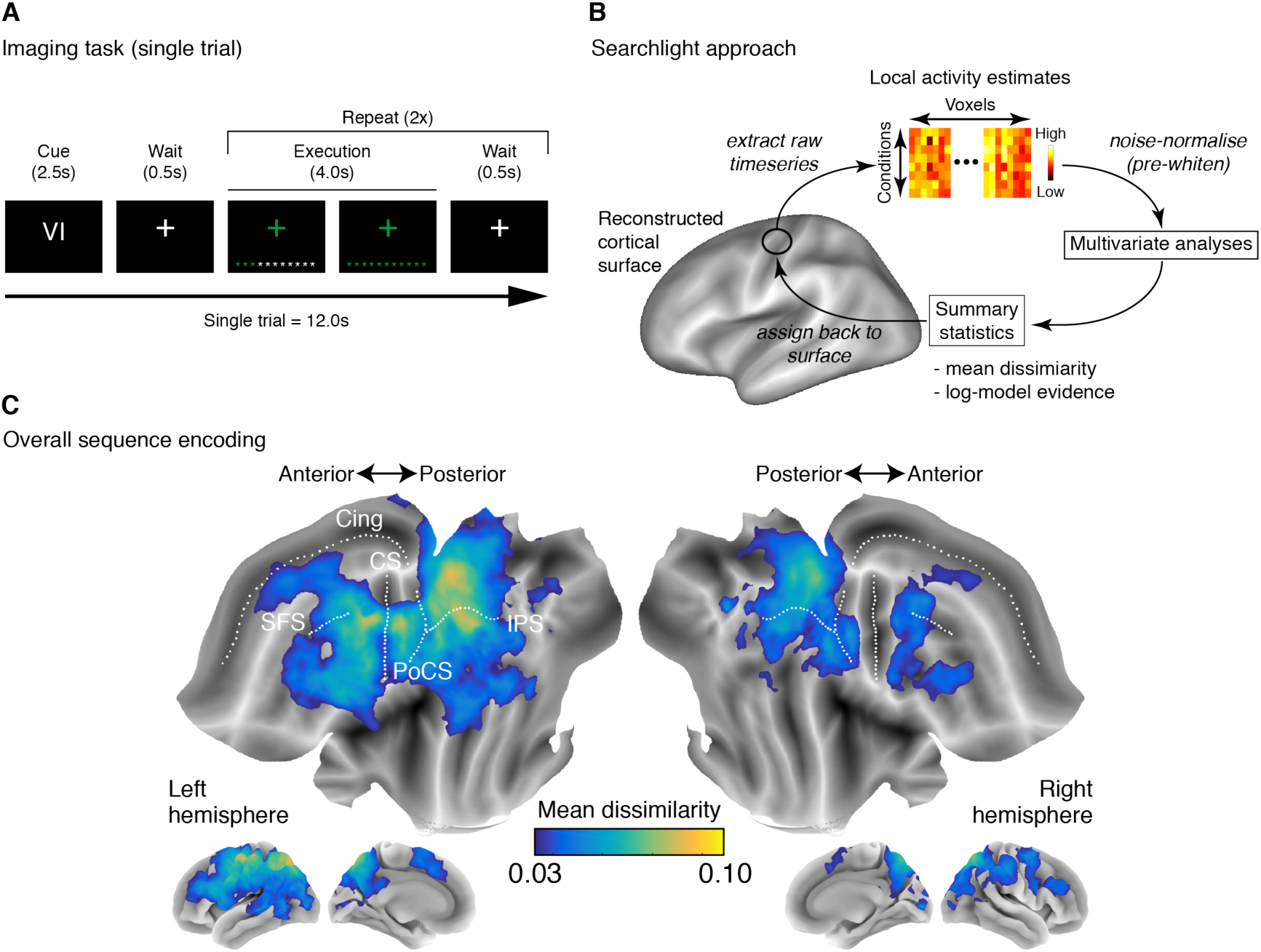
Overall sequence encoding on the flattened cortical surface. **(A)** Time course of a single trial in the fMRI session. A presentation of the visual cue (sequence I-VIII) was followed by two executions of that sequence. **(B)** Searchlight approach. We extracted the activity patterns for small circular areas (~22mm diameter) of the reconstructed cortical surface. The pre-whitened activity patterns were then used to calculate the crossnobis distance, an estimator for pattern separability, or submitted to PCM, which flexibly tests different representational models. The resulting statistics (mean distance across the sequences, or log-Bayes factor for each representational model) was then assigned back to the centre node of the region. **(C)** A group-averaged map of the strength of sequence encoding (average pair-wise distance between sequences) plotted on a flattened cortical map. Cing: cingulate sulcus, SFS: superior frontal sulcus, CS: central sulcus, PoCS: post central sulcus, IPS: intraparietal sulcus.

Figure 3C shows the resultant group searchlight map displayed onto the flattened cortical surface. Consistent with recent studies (Kornysheva and Diedrichsen, 2014; Wiestler and Diedrichsen, 2013; Wiestler et al., 2014; Yokoi et al., 2018), sequences were “encoded” over the wide area over the cortical surface, including M1, S1, PMd, and areas around the IPS. Notably, we also found significant encoding in SMA/pre-SMA, extending into the rostral cingulate zone (RCZ, Picard and Strick, 1996), although the signal from these areas was weak compared with other areas such as PMd or IPS (Kornysheva and Diedrichsen, 2014; Wiestler and Diedrichsen, 2013; Wiestler et al., 2014). While some areas showed bilateral representations, they were consistently strongest in the left, contra-lateral hemisphere. We also detected relatively strong sequence encoding in the left lateral prefrontal cortex and bilaterally in the precuneus (Fig. 3C).

### Determining the representational structure of sequence “encoding”

While Figure 3C tells us that we can decode the sequence identity from the activation pattern in this area, it does not answer whether these patterns differences are caused by neuronal populations that represent finger transitions, chunks, or entire sequences. In regions in which we observe significantly positive dissimilarities, we therefore looked in detail into the representational structure characterised by the representational dissimilarity matrix (or equivalently the second-moment matrix, Diedrichsen & Kriegeskorte, 2017) of the patterns (Fig. 4B).

**Figure 4.**
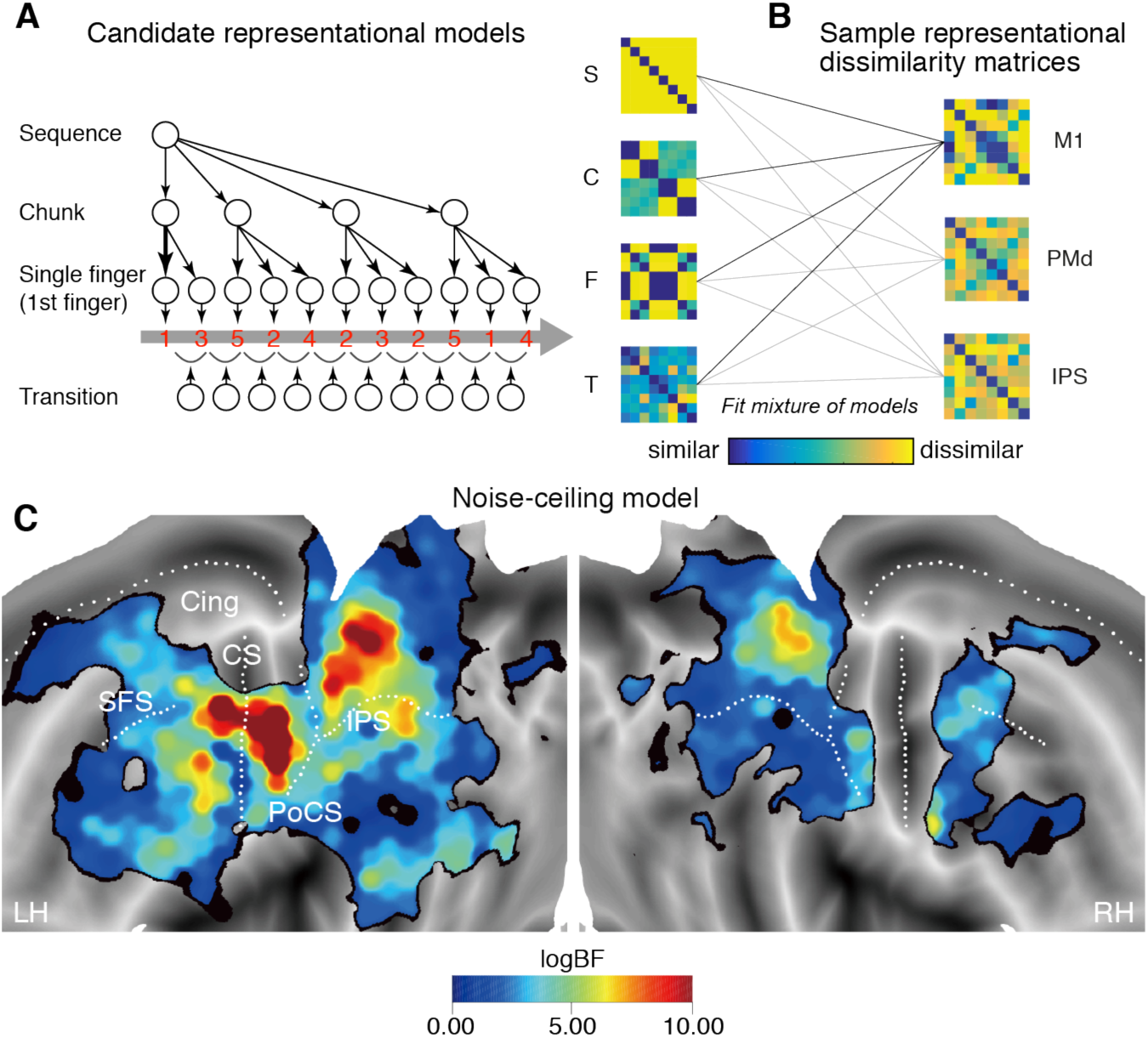
Fitting candidate representational models onto data. **(A)** Four candidate representational model components of an example sequence (red numbers). Each circle represents a hypothetical neural population; arrows between the circles represents descending commands to activate the units. For example, the sequence representation at the top sends ordered descending commands to the chunk representations, and each chunk representation sends commands to the single finger representation. Note that for the single-finger level, a unit for each single finger is shown multiple times. The non-hierarchical transition representation activates each press, based on the previous press. Given the sequences used (Fig. 1B), each representational component predicts a unique structure of the representational dissimilarity matrix (RDM). S: sequence, C: chunk, F: first-finger, T: transition. (**B**) Empirical RDMs for representative cortical regions. We fitted various combinations of the candidate models to explain observed representational structure at each cortical searchlight (see also Fig. 3b,c). **(C)** Group-level map of model evidence (log-Bayes factor) for the noise-ceiling over the null model. LH: left hemisphere, RH: right hemisphere, CS: central sulcus, SFS: superior frontal sulcus, PoCS: post-central sulcus, Cin: cingulate sulcus, and IPS: intraparietal sulcus. The logBF map was thresholded using a protected exceedance probability (PXP) of 0.75 (Rigoux et al., 2014; Rosa et al., 2010; Stephan et al., 2009).

Based on our behavioural results, we first considered three levels of hierarchical sequence representation (sequence, chunk, and single finger). At the highest level, we propose a sequence representation, with a unique neuronal activity pattern for each of the 8 trained sequences. As we assume that all sequences are equally strongly encoded, such a representation would predict that all possible pairwise distance are equal. A dedicated sequence representation would explain transitions between chunks are performed faster in trained than in novel sequences. At the next level, we have distinct neural activity patterns for each learned chunk. A region with a pure chunk representation would therefore transition during the sequence through the four activity states associated with the four chunks. The resultant RDM is therefore predicted by how many chunks different sequences have in common. For instance, sequence 1 and 2 consist of the same chunks in a different order (Fig. 1B) and are therefore predicted to elicit highly similar activity patterns. At the lowest hierarchical level, we considered representations of single fingers. As all sequences consisted of exactly the same presses arranged in a different order, a single-finger representation should lead to identical activity patterns for all sequences. In a recent study, however, we found that the activity pattern for a sequence is strongly determined by the first finger (Yokoi et al., 2018). Therefore, this representation predicts that sequences starting with the same finger should be very similar to each other (Fig. 4A). Thus, a single-finger representation would be detectable in this experiment as a first-finger representation. Additionally, we also considered that regions may show a distinct activity pattern for each finger transitions. The sequences were designed to maximally dissociate the predictions from these four models (sequence, chunk, first-finger, and transition).

With these candidate representational models, we applied pattern component modelling (PCM, Diedrichsen et al., 2011; Diedrichsen et al., 2017) to estimate the contribution of each candidate representational model to the observed activity patterns. PCM provides a direct and powerful Bayesian approach to test representational models (Diedrichsen and Kriegeskorte, 2017), and is functionally equivalent to RSA or encoding model approaches. Importantly, it provides a principled and flexible way to test for combinations of model components (or feature sets). We therefore used PCM to evaluate the likelihood of the data under all 16 possible combinations of the 4 candidate models within the general sequence “encoding” map (Fig. 3C, Fig. S1). The relative weight of each model component was fitted. Because different combination models had different number of free parameters, we used leave-one-subject-out cross-validation (Diedrichsen et al., 2017), fitting each model to all participants except one, and then evaluating the likelihood of the data from the left-out participant under the model (see Method for more detail).

Figure 4C shows the log-Bayes factor for a fully flexible “noise-ceiling” model, in which the predicted representation structure for each individual was the average representational structure for all the other participants (Walther et al., 2016) (see Method). Positive evidence for this model over the null model (no representation) simply indicates that the structure of representation was consistent across individuals. Almost all of the region tested survived the subsequent Bayesian group analysis (Rigoux et al., 2014; Rosa et al., 2010; Stephan et al., 2009) and its typical threshold (see Method). From the noise-ceiling map, we can clearly see that representational structure of the primary sensorimotor areas showed the highest inter-individual consistency being followed by areas around IPS and then frontal premotor areas (Fig. 4C).

Next, we examined whether these representations could be explained by our 4 candidate model components, or any combination of these. We therefore fitted all possible combination of components and then determined the model-averaged Bayes factor as a measure of evidence for the presence of each components in the context of the others (see methods).

### Representation of simple, elementary movements

Replicating our previous results (Yokoi et al., 2018), the representational structure in M1 and S1 was almost fully determined by the first finger in each sequence. The model-averaged log-Bayes factor revealed strong evidence for the first finger component (Fig. 5A). The logBF for the first-finger model was 1.26 ± 8.67 above the lower noise ceiling – which resulted in an insufficient protected exceedance probability (PXP) for the noise-ceiling model of 0.57. As the first finger press was executed with a similar force as all subsequent presses (t_14_ =0.42, p=0.34), this indicates that the same movement elicits more BOLD activity in M1 when it is executed in the beginning, rather than in the middle of a sequence. This simple model could fully explain the representational structure in these areas.

**Figure 5.**
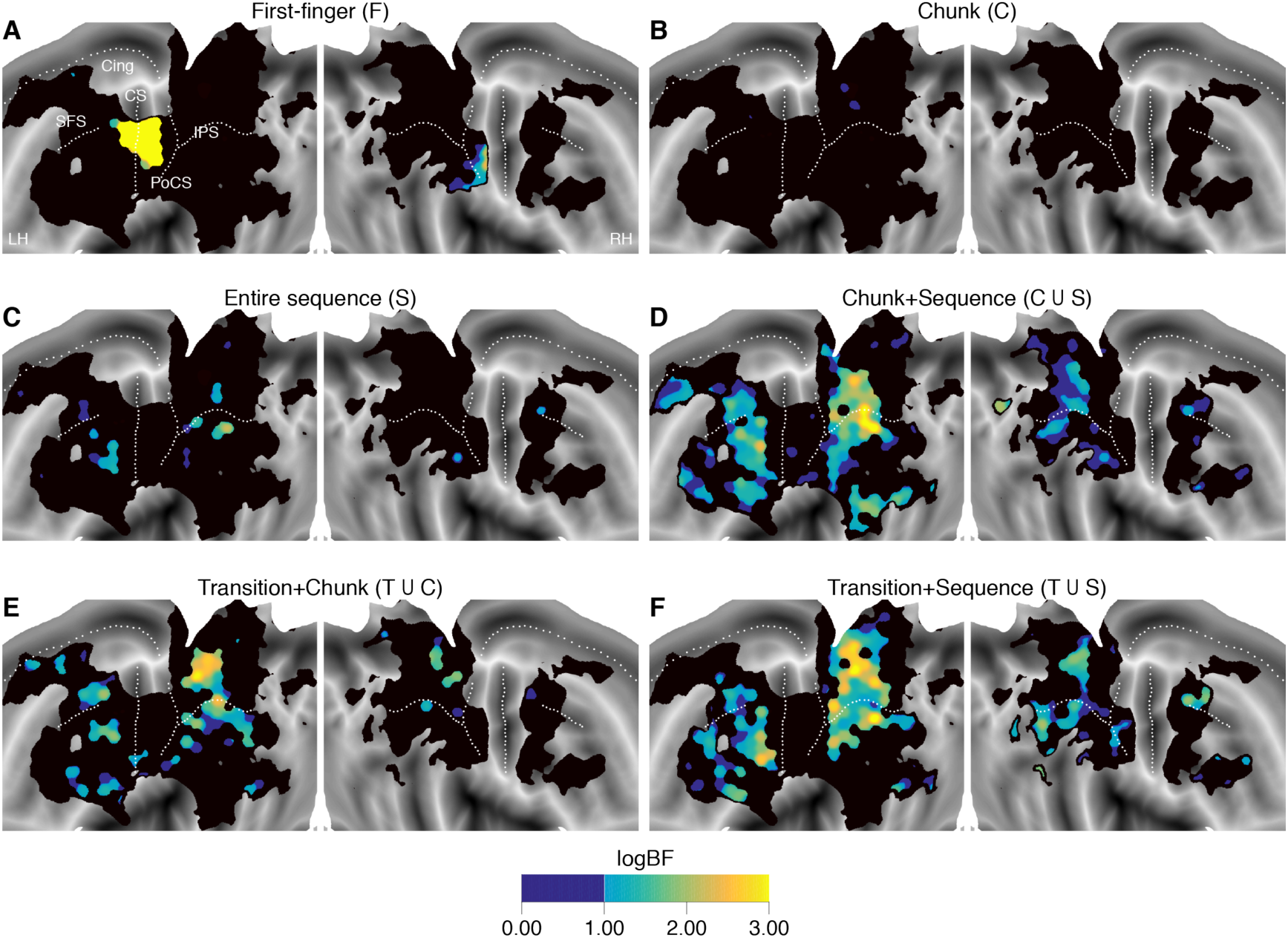
Bayesian model-averaging revealed elementary movements are robustly represented across individuals whereas higher-order representations are spatially overlapping. (A-F) Average log Bayes factor for first-finger (A), chunk (B), entire sequence (C), union of chunk and entire sequence models (D), union of transition and chunk models (E), and union of transition and sequence models (F) mapped onto flattened cortical surfaces. Each map was thresholded with a PXP of 0.75 (see Method). Areas with above-threshold PXP, but a logBF <1, are shown in dark blue.

### Representation of complex, higher-order movement hierarchy

In contrast to M1 and S1, the first finger model did not provide a good explanation for the representational structure in premotor and parietal areas (Fig. 5A). Model-averaging also revealed that there was only very weak or no evidence for either the chunk (Fig. 5B) or sequence model alone (Fig. 5C). Only when we assessed the average likelihood of the chunk, sequence, and sequence + chunk model (i.e., chunk ∪sequence), there was strong evidence in cortical regions along the left IPS and premotor areas (Fig. 5D). In these regions, the union of sequence and chunk models fitted the data significantly better than the lower noise-ceiling; logBF (union model vs. noise-ceiling) = 2.58 ± 3.29, PXP=0.952, for premotor cluster, and logBF =3.76 ± 4.83, PXP=0.85, for parietal cluster. We also found relatively strong evidence in left pre-SMA and more rostral medial regions, areas in the inferior frontal gyrus (including BA 44), right SPL and precuneus, as well as the right premotor cortex (Fig. 5D). The result suggests that chunk and sequence representations are not spatially segregated, but together can explain the structure of sequence representations in premotor and parietal cortices.

### Non-hierarchical movement representations

Similar to the chunk and entire sequence representations, there was almost no evidence for a representational structure that was explained by the 2-finger transition model alone (not shown in the figure). In contrast, we found firm behavioural evidence for learning of finger transitions (Fig. 2F). Therefore, we again assessed the evidence for the union of finger transition and other hierarchical (i.e., chunk or entire sequence) representations and found positive evidence similar regions where we found evidence for the hierarchical representations (Fig. 5E,F). For example, in left premotor and parietal cortices, the best fitting combination model included finger transitions. The result suggests that there is no spatial distinction between the neural populations for the hierarchical and the non-hierarchical movement representations.

### Model-free division of cortex according to the representational structure

The noise ceiling model (Fig. 4C) provided evidence for a systematic representational structure across participants in premotor and parietal areas, which could be modelled by a combination of chunk, sequence and transition representations (Fig. S2). However, there was no clear spatial separation of these representations. Rather, in most regions, at least 2 of the three components were necessary to account for the data, while others, such as bilateral precuneus and lateral prefrontal regions, were not well explained by any of these combinations. Therefore, it is likely that these areas have some unique way of representing sequences that was not captured in our candidate models. We therefore took a model-free approach to dissect cortical representation of sequences into discrete areas by applying an unsupervised clustering algorithm to the similarity (or “connectivity”) of the observed RDMs across the searchlight nodes within the areas that showed general encoding (Fig. 6A). The similarity between the resultant 10 clusters (Fig. 6B) could then be evaluated using an agglomerative hierarchical clustering algorithm which revealed that they formed several “families” (clusters 1-4, 5-7, and 8-10, Fig. 6C, for detail, see Method).

**Figure 6.**
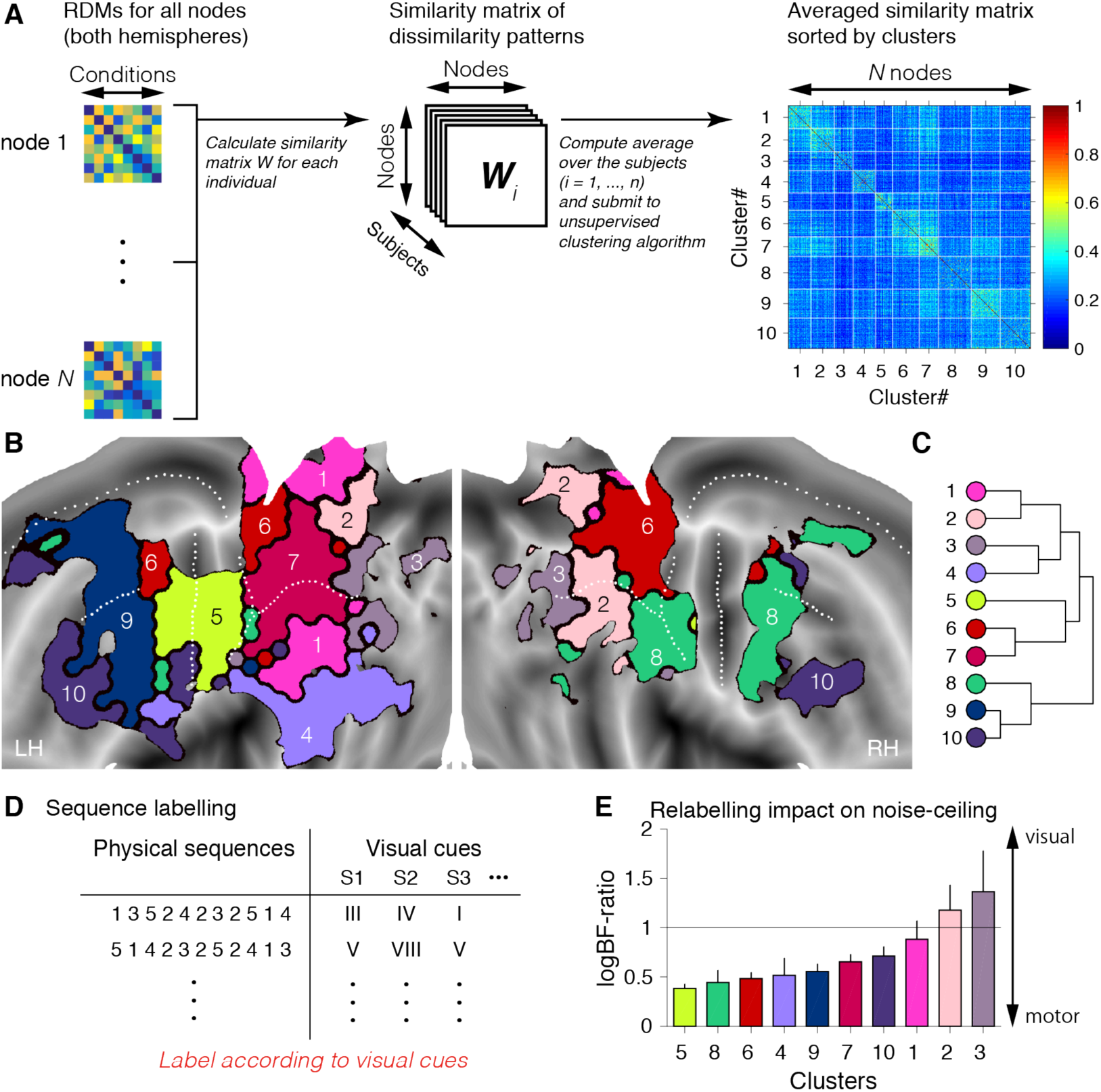
Parcellation of cortical motor sequence representations. **(A)** Schematic of the model-free clustering approach. **(B)** Resultant 10-cluster solution, mapped on the cortical surface. **(C)** Agglomerative clustering (Ward’s clustering) described relationships across the clusters. Colour similarity is directly related to cluster similarity **(D)** Condition relabelling according to actual visual cues each participant received during the task. Each participant received different visual cues for the same sequences. **(E)** A comparison between noise-ceilings obtained under a motor-based labelling and a visual cue-based labelling for each cluster. Vertical axis represents the logBF-ratio of noise-ceiling models (visual vs motor).

The resultant clusters revealed some features consistent with the model-based approach. Specifically, one cluster encompassing the left M1 and S1 captured the area that could be fully explained by the first-finger representation (cluster 5, Fig. S3). In premotor and parietal areas, the clustering was able to reveal features that could not be captured by the model-based approach alone. The approach showed that the regions with clear positive evidence for the mixture of higher-order representations (Fig. 5B,D,E,F) were sub-divided into several distinct clusters. For example, BA 5 and caudal PMd had a very similar representational structure (cluster 6), consistent with the strong anatomical connections shown between these areas (Kurata, 1991; Tomassini et al., 2007). A related cluster (cluster 7) accounted for the structure of the representation along the IPS. Further anterior, cluster 9 and 10 included left rostral PMd, SMA, and preSMA, and the dorsolateral prefrontal cortex, with cluster 8 encompassing the right PMd. When we applied the model-based PCM to these clusters, we found that they each showed a different mixing ratios of higher-order representations (Fig. S4).

### Cascade of information processing during sequence production

From the 4 remaining clusters, 1 and 2, as well as 3 and 4 formed one family (Fig. 6B). These areas, except for the cluster 4, showed only a very low inter-subject consistency in the PCM approach. One possible reason is the fact that we randomized the associations between the visual cues and the actual sequences across the participants. Because our “representational pracellation” approach is based on within-subject similarity of the RDMs, it is likely that these clusters captured either visual features of the presented cues (I, … VIII), or verbal features of internal rehearsal processes, rather than the features related to motor execution. Therefore, over the all clusters, we re-calculated the noise-ceiling after re-aligning the conditions across participants in terms of the visual cue identity, and compared it with the original noise-ceiling. The result demonstrated that the representational structure of clusters 2 and 3 was more consistent in “visual” than “motor” space (Fig. 6D,E). As expected, the cluster encompassing M1 and S1 was located at the most “motor” position, while the other clusters occupied positions that suggest a mixture of motor and perceptual / symbolic representations of the sequences.

In sum, the current results provide a set of novel insights into the cortical organisation of a sequential motor memory. First, we confirmed a clear hierarchy between M1 and premotor / parietal areas, with neural populations in primary sensory and motor area engaged in the generation of individual finger presses. In contrast, higher motor areas, such as PMd, SMA, and the parietal cortex showed true sequence dependency. While chunk, sequence, and transition representations together could account for these representations, no clear specialisation or hierarchical ordering emerged from this model-based analysis. Rather, most areas showed a mixture of representations between these regions. Our model-free approach provided the first insight into qualitatively different groups of representations. Finally, the relabelling analysis established the position of the clusters along the continuum between cue-oriented to more execution-related areas.

## Discussion

The current study provides the first direct evidence for a diverse set of neural representations of complex motor sequences in the human neocortex across frontal and parietal cortices. Our experimental approach was to impose a specific structure along which participants first build up a declarative knowledge of the sequences. We confirmed that this imposed structure was reflected in participants’ motor behaviour even after the need for memory recall was removed. We then could study the neural correlates of the sequence representation using both model-based and model-free multivariate fMRI analyses. The cortical clusters identified by model-free approach were further characterized in terms of stimulus-to-response gradient.

It has long been debated whether the consistent regularities in the timing of sequential behaviours reflects a hierarchical organization of sequence representations (Sakai et al., 2003), the associative learning of transition statistics (Verwey and Abrahamse, 2012), or merely arises from the biomechanical requirement at the specific finger transitions (Jimenez, 2008). Our experimental design with two groups of participants acquiring physically the same sequences through two cognitive routes provides clear behavioural evidence for chunking independent of the biomechanical property of those sequences (see also, Verwey and Dronkert, 1996). The results of follow-up experiment also indicate that the frequency of 2-finger transition independent of chunking influence performance. In the subsequent fMRI analysis, we confirmed that these two different types of representations coexist in premotor and parietal areas.

Our multivariate fMRI approach allowed a direct assessment of sequence representations. Many prior univariate studies on motor sequence learning have revealed experience-dependent activity changes in multiple brain regions, including DLPFC, M1, PM, SMA, IPS and precuneus (Doyon et al., 2002; Grafton et al., 1995; Honda et al., 1998; Kawashima et al., 1998; Penhune and Steele, 2012; Sadato et al., 1996; Sakai et al., 1998). A recent line of multivariate fMRI studies (Kornysheva and Diedrichsen, 2014; Nambu et al., 2015; Wiestler and Diedrichsen, 2013) has provided direct evidence that these previously reported regions represent some information about motor sequences. In the current study, we went one step further by characterizing this information in detail.

### Primary vs non-primary sensorimotor areas

Our data indicates that M1 and S1 only represent single finger movements, but no sequential information. The differences between the different sequences can be explained purely by the fact that the first finger had a stronger influence on the overall BOLD activity pattern than the remaining finger presses. This replicates our previous study (Yokoi et al., 2018) where we showed the same effect with balanced set of 6-digit sequences. In that study, we measured the patterns for each involved finger separately and demonstrated that each sequence was more similar to the pattern of the first finger. The paper also provides evidence that the first-finger effect has neuronal, rather than hemodynamic causes. One explanation for these effects is that M1 receives the greatest input drive from the higher motor areas and/or subcortical structures at the initiation of sequential movements to move from a resting to an activated state. Otherwise the neural state of M1 appears to be determined by the elementary movements only. The source of this signal may be neural populations in PFC or basal ganglia, which have been shown to fire most vigorously at the start of an action sequence (Fujii and Graybiel, 2003; Jin and Costa, 2010). In fact, we found evidence for the first-finger representation in basal ganglia ROIs, although the signal in these structures was much weaker compared to the cortical regions (Supplementary Information, Fig. S5).

In contrast, we found strong evidence for sequential representations outside M1/S1, most notably in PMd and parietal regions. Together, these results highlight the functionally different roles between primary and non-primary motor areas in production of voluntary sequential movements.

### Rethinking the hierarchical and non-hierarchical motor representations

Unexpectedly, premotor and parietal areas showed a mixture of different hierarchical and non-hierarchical sequence representations. This lack of spatial segregation between more complex representations might be simply due to that the underlying neural circuit for each representation is described in sub-voxel scale and hence could not be dissociated with the spatial resolution of current study. Alternatively, it may support the view that hierarchical behaviour itself does not necessary require hierarchically arranged neural architecture (Botvinick and Plaut, 2004; Yamashita and Tani, 2008). Our brain may possibly employ some unique way to organize hierarchical movement sequences without explicit architecture-level hierarchy (Koechlin and Jubault, 2006). Neural circuits in premotor and parietal cortices, which initially only transmit the selection signal from the “higher” cognitive areas, may self-organize as an “intermediate” layer into different types of representation in both “hierarchical” or “non-hierarchical” manner as movement sequences are repeated (Diedrichsen and Kornysheva, 2015). This may explain the co-existence of chunk-and transition-based motor skills found in our behavioural results. If so, the important next challenge is to understand the principles that govern the self-organisation of these networks.

An important concept in this context is the notion of “untangling” of the population response (Russo et al., 2018). To produce a temporally ordered sequence of neuronal signals, the generating neuronal region needs a representation of the sequential context, i.e., the neuronal state needs to be sufficiently different for movement A if it is followed by B, as compared to when A is followed by C. Similar neuronal states for movement A in these two contexts would lead to ‘tangling’ of the population response and the danger of confusion and would require substantial input to bring the neuronal dynamics on the correct path. Consistent with our findings, very recent results (Russo et al., 2018, personal communication) indicate that the neural state in M1 shows high tangling on the level of movements, whereas SMA appears to provide an untangled signal, where the neuronal state for the same movement depends on the sequential context. While this approach does currently not specify how this untangling is realized, it is conceivable that the neuronal population would develop an internal representation that looks like a mixture of a hierarchical on non-hierarchical organization.

### Representational parcellation of human neocortex during skilled sequence production

While many of existing attempts of cortical parcellation have been relying on correlations between time series across different brain regions, mostly during the rest (i.e., functional connectivity, Margulies et al., 2016; Yeo et al., 2011), our parcellation approach is unique in using correlations between representational structures (“representational parcellation”). The specific advantage of the current task-based representational parcellation approach is that in close combination with model-based representational fMRI analysis it allows us to make direct inference about the functional role for each network/cluster.

*Parietal family (Clusters 6 and 7):* Cluster 7 consisting of a large area along the left IPS was characterized by its strong evidence for higher-order representation (i.e., chunk + sequence, Fig. 5D). This fits well with the classical reports that lesions around the left IPS cause apraxia, a deficit arising from the loss of higher-order motor representations (Haaland et al., 2000). Additionally, these areas showed the second highest level of inter-subject consistency (Fig. 4C). Relabeling analysis according to the visual cue identity suggested that cluster 7 showed more perceptual / abstract representations than M1. Consistent with this idea, the areas along the IPS represent sequential movements in both intrinsic and extrinsic frames (Wiestler et al., 2014). Cluster 6 included both left caudal PMd and bilateral BA 5/precuneus, two brain regions with dense anatomical connections (Kurata, 1991; Tomassini et al., 2007), highlighting the critical role of caudal PMd as a bottleneck to translate the higher-order movement planning for the adjacent generating circuit (M1) (Dum and Strick, 2005; Ohbayashi et al., 2003). Consistently, a recent study has shown that inactivation of PMd in monkeys impaired the production of well-learned short motor sequences (Ohbayashi et al., 2016).

*Frontal family (Clusters 8-10):* The large complex of rostral PMd, PMv, SMA, and pre-SMA formed the cluster 9 and 8 on the left and the right hemispheres, respectively (Fig. 6B). More rostral prefrontal areas were classified into the cluster 10 bilaterally (Fig. 6B). While both showed significant model evidence for a mixture of hierarchical and non-hierarchical movement representations, the impact of cue-based relabelling on the noise-ceiling was the smaller in cluster 10 (Fig. 6E). These observations fit, at least partly, with the theory of rostro-caudal gradient of more abstract to concrete representation (e.g. Botvinick, 2008; Koechlin and Jubault, 2006). Thus, although highly intermixed, cluster 9 and 8 are more likely to represent concrete movement representations, such as chunks or transitions, whereas cluster 10 may contain more abstract representations involving the entire sequence. Such speculation is in line with the previous electrophysiological studies finding that neurons in pre-SMA, SMA and PMd specifically fire at the specific transition of successive movements (Ohbayashi et al., 2003; Ohbayashi et al., 2016; Tanji and Shima, 1994), and neurons in PFC, in turn, fire at the initiation of specific category of sequences (Shima et al., 2007).

*Peripheral family (Clusters 1-4):* The remaining clusters 1, 2, and 3 contained areas with relatively low level of model evidence and inter-subject representational consistency; bilateral precuneus, IPL (close to SMG), and the most posterior part of parietal regions, including pericarcaline sulcus (Figs. 4C and 6A). Activation of these areas has been reported to be related to sequence production and/or learning (Honda et al., 1998; Petit et al., 1996; Sakai et al., 1998). Interestingly, the spatial arrangement of the cluster 1 and 2 reflects the dense reciprocal connection between precuneus and other regions, including IPL and surrounding visual areas (Cavanna and Trimble, 2006; Margulies et al., 2009). Although highly speculative, the gradual reduction of relabelling effect in clusters 3, 2, and 1 might reflect the cascade of information processing. Together with the fact that precuneus is often activated during episodic memory retrieval (Cavanna and Trimble, 2006), these “networks” might reflect the participants’ retrieval of each sequence given the visual cues.

### Limitations & open issues

Our experiment was specifically designed to study how learned sequences are represented. Therefore, we are not able to track how the underlying representations change over the course of prolonged training. Furthermore, previous studies have demonstrated that sub-cortical structures, such as striatum, is one of the critical structures for the formation of movement chunks or habits (Graybiel, 1998; Graybiel and Grafton, 2015; Wymbs et al., 2012). Unfortunately, the spatial resolution of the current study (2.3 mm isotropic) was not optimal for applying MVPA to subcortical structures, resulting is relatively low noise-ceilings here (Supplementary Information, Fig. S5). An important future challenge is to understand the interplay between representations in sub-cortical and cortical structures in different stages of sequence learning.

## Conclusion

Our results provide evidence for a mixture of hierarchical and non-hierarchical sequence representations in premotor, prefrontal and parietal cortices. Using a model-free approach we were able to derive the first map of the sequence representations, suggesting differential roles for each cortical area. The next challenge is to understand how these representations are formed over the course of learning.

## Acknowledgements

This study was supported by a JSPS Postdoctoral Fellowship (#15J03233) to AY, a James S. McDonnell Foundation Scholar award, a NSERC Discovery Grant (RGPIN-2016-04890), and a Platform Support Grant from Brain Canada and the Canada First Research Excellence Fund (BrainsCAN) to JD. We thank Elizabeth Bamber and Emily Thomas for the assistance in data collection, Naveed Ejaz, Uri Hertz for helpful discussions, Eva Berlot for comments on the manuscript.

## Author Contributions

A.Y. and J.D. designed the experiment. A.Y. collected and analyzed the data. A.Y. and J.D. wrote the manuscript.

## Declaration of Interests

The authors declare no conflict of interests.

## Materials & Methods

### Participants

All experimental procedures were approved by local ethics committee at the University College London (London, UK). We recruited 23 healthy, right-handed, neurologically healthy volunteers, who participated in the study after providing written informed consent. None of the participants was a professional musician. Of these, 8 participants were excluded, as they did not meet the performance criterion necessary to go on to the imaging session and post-test. The remaining 15 participants went through the imaging session, and the subsequent post-test. Of these 15 remaining participants, the data from 3 participants were excluded, as they failed to achieve sufficient behavioural performance during scanning (57% correct vs. 81% correct for all other subjects). As a result, only the data from the remaining 12 participants (5 females, 7 males, age: 23±4) was submitted to analysis. These participants reported 5.8±3.8 years of practice with musical instruments (e.g., piano, guitar, violin, etc.).

### Apparatus

We used a custom-built five-finger keyboard device (Fig 1A). The keys of the device were immobile and equipped with force transducers that could measure isometric finger forces (Wiestler and Diedrichsen, 2013; Yokoi et al., 2017). The analog signals were passed through a penetration panel in the magnet room to avoid radio-frequency leakage. The signals were then low-pass filtered, amplified, digitized, and sent to PC for online task control and data recording. The forces were recorded at 200 Hz. During the training sessions, the participants placed only their right hand on the keyboard to perform the task, while in the scanner the fingers of their left hand were placed on a mirror-symmetric device to monitor potential implicit mirror movement (Diedrichsen et al., 2013).

### Procedure (behavioural)

#### Sequences production task

We employed a discrete sequence production (DSP) task, in which participants were asked to produce a specific sequence of key presses as fast and accurate as possible. Over the course of 5-6 days (∼2 hours per each day), participants learned to produce 8 different sequences consisting of 11 presses from the memory as quickly as possible. All the sequences were matched with the number of finger presses used; 2 presses with thumb, middle, ring, and little fingers, and 3 presses with index finger, respectively. A finger press was detected when the force crossed a threshold of 3 N and a release was detected when it fell below the threshold. To successfully complete a finger press, the pressed finger needed to be pressed, while all other finger needed to be released.

During the imaging task, a central fixation cross was presented. Each trial started with a 2.5s of the presentation of a visual cue (roman numerals I-VIII) that indicated the sequence to be executed. The cue presentation was followed by 0.5s of interval. Then, the fixation cross turned green, and 11 asterisks were presented, triggering the subject to produce the sequence (Fig. 3A). For each correct press the corresponding asterisks turned green - for each incorrect press red. Participants were instructed to complete the sequence even if an error has occurred. After each execution, feedback was given (during the ITI) by the colour of fixation cross (white: correct, red: one or more presses were incorrect, and blue: unfinished, but presses were correct).

#### Behavioural training

In order to ensure consistent and stable chunk structure across individuals, we deliberately imposed chunk structure by manipulating how the participants build up their explicit memory of the sequences. In brief, we initially trained the participants with single “chunks” (Fig. 1B) until they could produce these chunks from memory, and then trained them with sequences that consisted of these chunks. Eight different chunks of 2 or 3 presses (cued by alphabets) were organized into 8 different sequences, every one of which consists of combination of 4 chunks (Fig. 1A). The associations between chunks and chunk cues (A, B, ~, H), and between sequences and sequence cues (I, II, ~, VIII) were randomised across participants. To dissociate the influence of the explicit training from subsequent biomechanical optimisation of the sequence, we assigned participants randomly to one of two groups, which were trained with different set of chunks. Five sequences were physically identical across the two groups (i.e., the same order of finger presses chunked differently, Fig. 1B). The sequences were designed to maximize the difference in the prediction of the different representations models (see below).

Training consisted of 5 days before the imaging session. On day 1 and early blocks of day 2, subjects were specifically trained with individual chunks. We alternated cued trials in which the chunk cue (A-H) was presented together with the required digits, and uncued trials where only the chunk cue was presented. In each block, each chunk type was repeated for three times. Participants received a total of 720 trials of chunk training. Starting on day 2, participants practices entire movement sequences. On cued trials, the sequence cue (I-VIII) and chunk cues were presented (Fig 1C), but no longer with finger cues. On uncued trials, participants needed to retrieve the entire sequence from memory. In each block, each sequence type was repeated for three times. They received a total of 1512 trials of sequence training.

On the fifth day of the training session, after the practice session, the participants practiced the task in a supine position on a mock MRI scanner bed. For the half of the participants, we added the 6th day of additional familiarization session to ensure that they could correctly produce the sequences within 4 seconds. They were familiarized with the actual task in the scanner during on average 10±8 blocks of the familiarization session.

#### Post-test session

To confirm that the cognitively imposed chunk structures actually influenced participants’ motor representations, we conducted a post-test session within 1 week after the imaging session (1±0.6 days). In the session, all the digits were presented on the screen to release the participants from the necessity to recall any sequence from memory. No sequence cues were provided. Additionally, we assessed the generalization of learned chunks to unlearned sequences by additionally introducing 3 new sets of sequences (Fig. 2D): New: completely novel sequences which did not contain any of the trained chunks; Chunk: sequences composed of trained chunks in untrained order; and Chunk + New: novel sequences that contained two learned chunks at random positions in the sequence. Each category except for Chunk, which had 16 different sequences, had 8 different sequences, resulting in totally 40 sequences. Sequences were executed 4 times in a row. The order of sequences was randomized and all sequences were repeated for 4 times (16 executions per a sequence). The resultant 640 executions were divided into 16 blocks.

#### Imaging session: Behaviour

During the imaging session, the participants were placed on the scanner bed with their knees slightly bent and supported by a wedge-shaped cushion. The two keyboards were tied together with plastic screws and stabilised on the participants’ lap with foam pads. Visual stimuli were presented on a back-projecting screen, and participants viewed the screen through the mirror mounted above the head coil. The presentation of the sequence cue (2.5 s), was followed by two execution phases for the same sequence (4 s for each). After this, the next trial started after an ITI of 0.5 s (Fig. 3A). The order of 8 sequences were randomized and each sequence had 3 trials (with 2 executions each) within each functional imaging run. Each run also contained 4 randomly interspersed rest phases (12 s). The number of correct trials for the run was presented in the screen at the end of the run. Each functional run lasted about 7 min and 9 runs per participant were conducted. Short breaks (up to a few minutes) were interleaved on the participants’ request.

#### Imaging data acquisition

Imaging sessions were conducted on a Siemens Trio 3T scanner system with a 32-channel head coil at the Welcome Trust Centre for Neuroimaging (London, United Kingdom). B0-field maps were acquired at the beginning of the session to correct for inhomogeneities of the magnetic field (Hutton et al., 2002). Functional images were acquired for 9 runs of 135 volumes each, using a 2-D echo-planer imaging sequence (TR = 2.72 sec, in-plane acceleration factor = 2, resolution = 2.3mm isotropic with 0.3 mm gap between each slice, and 32 slices interleaved). The slices were acquired in an axial orientation and covered the dorsal aspects of the brain, including most of the frontal, parietal, occipital lobes, and basal ganglia. The ventral aspects of the frontal and temporal lobes, brainstem, and the cerebellum were not scanner. The first 5 volumes of each run were discarded to ensure stable magnetization. A T1-weighted anatomical image was obtained using MPRAGE sequence (1mm isotropic resolution).

#### Behavioural data Analysis

Recorded force data were analyzed offline. Reaction time (RT) from the go cue, movement time (MT) starting from first finger press to the last finger release, inter-press intervals (IPIs), and the number of incorrect presses at each execution were calculated. An IPI was defined as the interval between two consecutive press onsets. We obtained similar result when using the interval between two consecutive peak force times as a measure of IPI.

#### Linear IPI modelling

To assess the contribution of imposed chunk structure and other effects, we ran a linear regression analysis on the IPI data of the follow-up session. We treated the individual IPIs for all trials as a single data vector. We then built linear models to explain the variation in the IPIs. These models consisted of all possible combinations of the following components: 1) transition: the frequency of the specific digit transition (25 total) in training, the sign of the regressor was negative, so that higher frequencies would predict lower IPIs; 2) chunk: whether the IPI was within a chunk (coded as −1) or not (0); 3) sequence: whether the between-chunk transition was trained (−1) or not (0); 4) biomechanical difficulty: the mean IPI of that particular finger transition in a control experiment that tested for the execution speed of all possible 2-and 3-finger transitions, as will be reported in a separate paper, and 5) post-error slowing: whether the proceeding press was incorrect (1) or correct (0). All models also included an intercept. All components, except for the intercept part, were then z-standardized before entering them into the regression analysis. For each model tested, we calculated AICc. For Fig 2F, the regression coefficients for the models were averaged by using the following Akaike-weight;

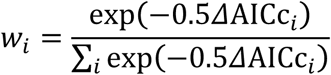

,where *w*_*𝒾*_ is the Akaike-weight for the model *i*, ΔAICc_*𝒾*_ is the difference in AICc between the model *i* and the best model. Such model averaging gives better prediction accuracy than using the best model alone (Burnham and Anderson, 2004). For averaging, the regression weights were treated as zero when a model combination did not contain those terms. The resultant model-averaged regression weights separately calculated for the participants were then submitted to the group statistical test.

### Imaging data analysis

#### Preprocessing and first-level model

Functional imaging data were pre-processed using SPM 8 (http://www.fil.ion.ucl.ac.uk/spm/). Functional images were first slice-time-corrected, motion corrected, and then co-registered to the individual anatomical image. We also corrected for B0 inhomogeneity by using field map images when correcting the head motion. The data were then submitted to a 1st-level GLM to estimate the size of the evoked activity for each sequence in each run. We used the standard high-pass filtering with a cut-off frequency of 128s before GLM estimation. We applied robust-weighted least square estimation (Diedrichsen and Shadmehr, 2005) to reduce the effect of any motion-induced artefact.

Each trial was modelled as a boxcar function, starting at the presentation of the go-cue with a length of 7.5 s for each execution. The boxcar function was then convolved with a standard hemodynamic response function. In the GLM used for subsequent analysis we included the activity of both correct and incorrect trials in the analysis. This was justified by two reasons. First, even if participants made a mistake, they were instructed to complete the sequence. This often happened automatically, as a substantial number of errors arose from omissions, in which the participants did not apply enough force to have the press registered. Secondly, given that each trial consisted of two executions of the sequences, many trials consisted of a correct and incorrect execution. Given the low temporal resolution of fMRI, we had little power to resolve this. An alternative analysis in which we excluded error trials from the activity estimation yielded similar, albeit noisier results.

#### Surface-based analyses

Our primary focus of analysis was cortical surface. We first reconstructed individual cortical surfaces (i.e., the pial and white-grey matter surfaces) from the anatomical image by using Freesurfer software (Fischl et al., 1999). The reconstructed cortical surfaces were then registered to a common symmetrical template (“fsaverage_sym”) (Greve et al., 2013). Subsequently, we defined the surface-based searchlight (Oosterhof et al., 2011) as small circular patches that contains 160 voxels (approximately 11 mm radius) centred on each node which was defined on the reconstructed cortical surface. The activity patterns of these 160 voxels for each centre were submitted to the multivariate analysis, and the result from each searchlight was re-assigned to the centre (for detail, see *Multivariate fMRI analyses*). The overall result was then used to restrict the regions of interest for further analysis.

For detailed testing of representational models, we defined a discrete searchlight parcellation. In contrast to the continuous searchlight map, we aimed to define a reduced set of only partly overlapping searchlights. This was done for computational efficiency for model testing, as well as for the model-free clustering approach. The searchlight centres were chosen only within regions in which the continuous searchlight analysis showed an average pattern distance across the sequence conditions greater than 0.03 (Fig. 3C). Within this region, we defined hexagonally-arranged searchlight centres on the flattened cortical surface coordinate for both hemispheres (using a 7 mm of spacing). We then defined a circular area around each surface nodes, such that each searchlight contained 150 voxels. This definition resulted in 465 (302 for the left hemisphere) partly overlapping tiles on the cortical surface (Fig. S1).

#### Multivariate fMRI analyses overview

For each of the defined searchlight, the beta-weights for each sequence type for each imaging run were extracted. The resultant beta-weights across voxels were then spatially pre-whitened by using multivariate noise-normalization with a regularized estimate of the spatial noise-covariance matrix (Walther et al., 2016). As a result, the activity estimates across voxels became approximately uncorrelated with respect to the noise (Diedrichsen and Kriegeskorte, 2017). We first took an RSA-approach to restrict the regions where the sequences are encoded. We then applied PCM to the regions with substantial sequence encoding to assess detailed content of sequence encoding.

The key quantity for both of these representational analysis techniques is the second moment (**G**) of the patterns for the sequences. The second-moment matrix is a covariance matrix, where the mean activity for each condition (across voxels) is not subtracted out (Diedrichsen and Kriegeskorte, 2017; Diedrichsen et al., 2017). Thus, when two patterns for sequence *i* and sequence *j* are similar to each other, the corresponding *i*,*j*^*th*^ element of **G** has a high value. For RSA, we computed the cross-validated estimate of **G** and derived from this cross-validated pattern distances. For PCM, we explicitly modelled the structure of **G** and then evaluated the likelihood of the data under these different models.

#### Overall sequence encoding

To define the cortical region which reliably encode different sequences, we assessed the discriminability of the elicited activity patterns on the surface-based searchlight. For this purpose, we first calculated a cross-validated estimate of the second moment matrix **G** as,

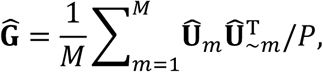

where *M* is the total number of imaging runs, *p* is the number of voxels within a searchlight, Û _*m*_ is estimated pre-whitened activity pattern for the *m*-th imaging run, and Û_*~m*_ is the estimate of the pattern independent of the *m*-th imaging run. Both of Û*~* _*m*_ and Û~ _*m*_ have size of 8×p. We then computed a cross-validated distance estimate from.Ĝ The squared cross-validated Mahalanobis distance estimator (crossnobis for short, Diedrichsen et al., 2016), between activity estimates for sequence 1, **u**_1_, and for sequence 2, **u**_2_, can be calculated as,

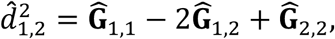

where Ĝ_*𝒾 ј*_ is the *i*,*j*^*th*^ element of **Ĝ**. We calculated the mean of all pair-wise crossnobis distance estimators across the sequences at each searchlight (Fig. 4). The crossnobis estimator is unbiased – meaning it can be used to directly test whether a distance is larger than zero. Finding consistently positive distance estimates therefor implies that the two condition activity patterns differ from each other more than expected by chance

#### Model-based approach

The above analysis is sensitive to any possible differences between the patterns associated with the different sequences. To dissect different forms of sequence representation, we used pattern component modelling (PCM) that allows to model the covariance structure (second moment matrix) across the activity patterns according to different representational hypotheses (Diedrichsen and Kriegeskorte, 2017; Diedrichsen et al., 2011; Diedrichsen et al., 2017; Yokoi et al., 2018). For our experiment, we defined the following five representational model components, including a null-model which predicts no difference across the sequences.

#### Sequence model

This model component assumes that each sequence is associated with unique activity pattern with common variance,

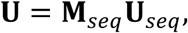

where the weighting matrix **M**_*seq*_ is an identity matrix (i.e., **M**_*seq*_ *=***_*I*_**_*8*_) and the pattern **U**_seq_ is uncorrelated (i.e., 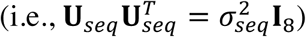. Therefore, the predicted second moment matrix has the simple form of,

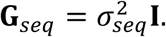

#### Chunk model

The chunk model assumes that the activity for each sequence is a combination of activities associated with the chunks it contains,

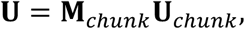

where the weighting matrix **M***chunk* specifies the membership of chunks used in each sequence (e.g., 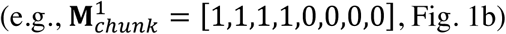 Fig. 1b), and the pattern **U**_*chunk*_ is also assumed to be uncorrelated (i.e.,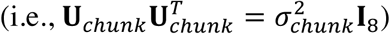 Therefore, the predicted second moment matrix has the specific form that reflects the composition of sequences in terms of chunks, 

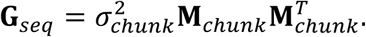

#### 1st-finger model

We have previously shown that differences in the sequence-specific activity pattern of M1 and S1 can we well explained by the fact that the first finger press shows a particularly strong activation compared to the subsequent finger presses (Yokoi et al., 2018). Because each sequence contained each finger equally often (and because the peak force of all finger presses was approximately the same), we can assume that the only thing that would differentiate these sequences in a region that only encodes single-finger movements, is which finger starts the sequence. The first-finger model therefore characterizes the part of the activity pattern that is different between sequences as a scaled version of the pattern associated with the first finger,

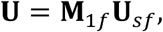

where the 8×5 weighting matrix **M**_1*f*_ has a scaler σ_1*f*_ at the column corresponds to the starting finger of each sequence (row), and the **U**_s*f*_ is the activity patterns associated with single finger presses. As we did not measure the single finger activity **U**_s*f*_ for the current experiment, we utilized the fact that the second moment of single finger patterns **G**_s*f*_ in M1 and S1 is well-characterised by the natural statistics of hand usage (Ejaz et al., 2015). The predicted second moment matrix is therefore,

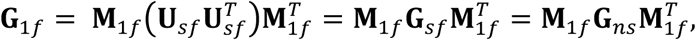

where **G**_*ns*_ is a second moment matrix predicted from the natural statistic of hand movement (Ejaz et al., 2015). Replacing G_*ns*_ with the second moment matrices for single finger representation that was derived from an independent experiment (Yokoi et al., 2018) did not affect the results. The second moment matrix predicted by this model therefore reflects which sequences share the first finger, and how similar the respective finger representations are to each other.

#### Two-finger transition model

As an alternative to the hierarchical representation of sequences, this transition model assumes that every sequence is represented as a combination of specific transitions that are defined continuously between each pair of movements (e.g., sequence 1-3-2-4-1-5 is composed of transitions 1-3, 3-2, 2-4, 4-1, and 1-5). Similar to the previous models, the activity patterns are modelled as,

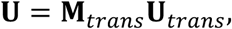

where the weighting matrix **M**_trans_ describes which of specific transitions occur in each sequence. The pattern **U**_trans_ are assumed to be uncorrelated with each other, but the strength that a transition *i* is represented was assumed to be proportional to the frequency of occurrence in training f_*i*_ (i.e.,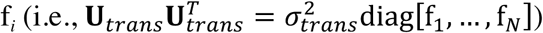. The predicted second moment matrix is therefore,

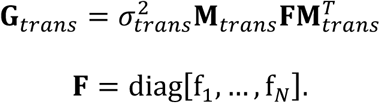

Dropping the assumption of training-dependent strength of transition representations (and hence setting **F** to the identity matrix) did not change the results qualitatively.

#### Null-model

As a baseline to evaluate each model, we defined a null-model that hypothesized no difference between any of the sequence patterns. For this, the hypothesized second moment matrix was

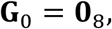

Where **0**_8_ is a 8×8 matrix whose elements are all 0.

#### Noise-ceiling model

In addition to the above models, we also fitted a fully-flexible noise-ceiling model to assess the maximally explainable information shared across individual. Here, we used a naïve noise-ceiling model that uses the empirical, cross-validated estimate of the second moment matrix as the predicted second moment matrix;

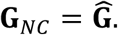

We fitted this noise-ceiling model separately for the two groups, as these groups of participants practiced partly different set of sequences with different chunking structure. We then combined the evidence for both groups for evaluation.

#### Model design

The prediction of each model was determined for each group separately, as they differed in both sequences and chunks. Because the two groups had 5 sequences that were identical, but were chunked differently, difference in the representational structure across these shared sequences could be specifically attributed to the difference at the level of the chunk representation.

Importantly, the activity estimates for each trial contained trials that either contained an error, or were not completed (see First-level modelling). We included these trials, because even for these incorrect trials, most of the sequence was produced correctly. To account for the fact that the beginning of the sequence was more often executed than the end of the sequence, we weighted each element of the representation (e.g., the first, second, third and fourth chunks or the first to tenth transitions) by the relative %-correct with which this element was produced. For example, the %-correct of chunks were on average 96, 94, 86, and 82%, hence each chunk element in the PCM model was weighted accordingly, which slightly changed the structure of corresponding weighting matrix.

#### Model evaluation

First, we fitted the all 16 combinations of above models (i.e., first-finger, 2-finger transition, chunk, and entire sequence models). As each of the combination models had different number of free parameters, i.e., combination weights, we evaluated the models using leave-one-subject-out cross-validation to prevent overfitting (Diedrichsen et al., 2017). Because the combination weights should be the same for the same search region (i.e., searchlight) across the two groups of participants, they were constrained to be the same across the two groups. We then used the resultant cross-validated log-likelihood (ℒ_*m*_) for each model *m* as an estimate of the model evidence. The log Bayes factor (BF) of model A over model B can then be calculatedas

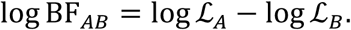

The evidence for each of the model components in the context of all other components can then be calculated (as for the behavioral analysis) using Bayesian model averaging. The log BF for each component is then the sum of the posterior probabilities for the models that contained the component (c=1) versus the sum of the posterior probability for the models that did not (c=0) (Burnham and Anderson, 2004; Shen, 2018).

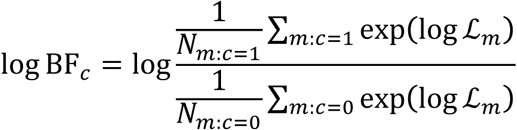

To assess the evidence for the union of two components models, we summed the posterior probability of all models that contained either or both of these components. For visualization, except for the Figure 4C, we employed a typical threshold for the logBF (logBF=1, Kass and Raftery, 1995).

#### Model-free approach

As a complement of our model-based approach, we also applied a model-free clustering of cortical surface regions based on their representational structure. As an input data, we used the crossnobis distance estimator (see *Overall sequence encoding*) for all the 28 pairs across the 8 sequences calculated at each searchlight (i.e., input data is an *N*-by-28 matrix, where *N* is the number of searchlight centers). We calculated an *N*-by-*N* similarity matrix across the searchlight centers for each participant by first calculating correlation distance d=(1-r) across the nodes. We chose the correlation distance in order to emphasize the profile of dissimilarity, rather than the magnitude. The resultant correlation distances were then transformed into similarity using a Gaussian similarity transformation;

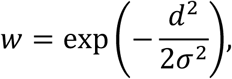

where *w* is the similarity, *d* the correlation distance, and σ the width of Gaussian. The width was determined individually as lower 5 percentile value of the correlation distances. We then applied the spectral clustering algorithm (Von Luxburg, 2007) to the group-averaged-similarity matrix with Jordan-Weiss normalization (Ng et al., 2002). To assess the similarity across the estimated clusters, we further applied agglomerative hierarchical clustering (Ward’s method) to the resultant cluster-average of Laplacian eigenvectors.

### Inferential Statistics

#### Behavioural data analysis

To compare within-and between-chunk intervals, we used paired t-test. To test group-specificity of press-interval patterns, pairwise correlation coefficients (Pearson’s *r*) across the subjects (within-and across-group) were first z-transformed and then submitted to a two-sample *t*-test. The individual AIC model-averaged regression weights were tested by one-sample *t*-test.

#### Imaging analysis

We applied a group-level Bayesian analysis (Rigoux et al., 2014; Rosa et al., 2010; Stephan et al., 2009) on the log-Bayes factor data of the participants (*spm*_*bms()* function implemented in the SPM 12). Group-log-Bayes factor map was thresholded in terms of the protected exceedance probability (PXP) by 0.75 which is the posterior probability of a model being greater than 0.5 (i.e., probability that an effect is present more than a half of subjects) (Rosa et al., 2010). All the statistical analyses were performed on MATLAB (Mathworks, Inc.).

## References

Baldauf, D., Cui, H., and Andersen, R.A. (2008). The posterior parietal cortex encodes in parallel both goals for double-reach sequences. Journal of Neuroscience 28, 10081–10089.

Ban, H., and Welchman, A.E. (2015). fMRI analysis-by-synthesis reveals a dorsal hierarchy that extracts surface slant. Journal of Neuroscience 35, 9823–9835.

Botvinick, M., and Plaut, D.C. (2004). Doing without schema hierarchies: a recurrent connectionist approach to normal and impaired routine sequential action. Psychol Rev 111, 395–429.

Botvinick, M.M. (2008). Hierarchical models of behavior and prefrontal function. Trends in cognitive sciences 12, 201–208.

Botvinick, M.M., Braver, T.S., Barch, D.M., Carter, C.S., and Cohen, J.D. (2001). Conflict monitoring and cognitive control. Psychol Rev 108, 624–652.

Burnham, K.P., and Anderson, D.R. (2004). Multimodel inference - understanding AIC and BIC in model selection. Sociol Method Res 33, 261–304.

Cavanna, A.E., and Trimble, M.R. (2006). The precuneus: a review of its functional anatomy and behavioural correlates. Brain 129, 564–583.

Chikazoe, J., Lee, D.H., Kriegeskorte, N., and Anderson, A.K. (2014). Population coding of affect across stimuli, modalities and individuals. Nature neuroscience 17, 1114.

Diedrichsen, J., and Kornysheva, K. (2015). Motor skill learning between selection and execution. Trends in cognitive sciences 19, 227–233.

Diedrichsen, J., and Kriegeskorte, N. (2017). Representational models: A common framework for understanding encoding, pattern-component, and representational-similarity analysis. PLoS Comput Biol 13, e1005508.

Diedrichsen, J., Ridgway, G.R., Friston, K.J., and Wiestler, T. (2011). Comparing the similarity and spatial structure of neural representations: a pattern-component model. Neuroimage 55, 1665–1678.

Diedrichsen, J., and Shadmehr, R. (2005). Detecting and adjusting for artifacts in fMRI time series data. Neuroimage 27, 624–634.

Diedrichsen, J., Wiestler, T., and Krakauer, J.W. (2013). Two distinct ipsilateral cortical representations for individuated finger movements. Cereb Cortex 23, 1362-1377.

Diedrichsen, J., Yokoi, A., and Arbuckle, S.A. (2017). Pattern component modeling: A flexible approach for understanding the representational structure of brain activity patterns. Neuroimage.

Diedrichsen, J., Zareamoghaddam, H., and Provost, S. (2016). The distribution of crossvalidated mahalanobis distances. ArXiv.

Doyon, J., Song, A.W., Karni, A., Lalonde, F., Adams, M.M., and Ungerleider, L.G. (2002). Experience-dependent changes in cerebellar contributions to motor sequence learning. Proc Natl Acad Sci U S A 99, 1017–1022.

Dum, R.P., and Strick, P.L. (2005). Frontal lobe inputs to the digit representations of the motor areas on the lateral surface of the hemisphere. The Journal of neuroscience : the official journal of the Society for Neuroscience 25, 1375–1386.

Ejaz, N., Hamada, M., and Diedrichsen, J. (2015). Hand use predicts the structure of representations in sensorimotor cortex. Nature neuroscience 18, 1034–1040.

Fischl, B., Sereno, M.I., Tootell, R.B.H., and Dale, A.M. (1999). High-resolution intersubject averaging and a coordinate system for the cortical surface. Human brain mapping 8, 272–284.

Fujii, N., and Graybiel, A.M. (2003). Representation of action sequence boundaries by macaque prefrontal cortical neurons. Science 301, 1246–1249.

Grafton, S.T., Hazeltine, E., and Ivry, R. (1995). Functional mapping of sequence learning in normal humans. Journal of Cognitive Neuroscience 7, 497–510.

Graybiel, A.M. (1998). The basal ganglia and chunking of action repertoires. Neurobiol Learn Mem 70, 119–136.

Graybiel, A.M., and Grafton, S.T. (2015). The striatum: where skills and habits meet. Cold Spring Harb Perspect Biol 7, a021691.

Greve, D.N., Van der Haegen, L., Cai, Q., Stufflebeam, S., Sabuncu, M.R., Fischl, B., and Brysbaert, M. (2013). A surface-based analysis of language lateralization and cortical asymmetry. J Cogn Neurosci 25, 1477–1492.

Haaland, K.Y., Harrington, D.L., and Knight, R.T. (2000). Neural representations of skilled movement. Brain 123 (Pt 11), 2306–2313.

Hikosaka, O., Nakahara, H., Rand, M.K., Sakai, K., Lu, X., Nakamura, K., Miyachi, S., and Doya, K. (1999). Parallel neural networks for learning sequential procedures. Trends Neurosci 22, 464–471.

Honda, M., Deiber, M.P., Ibáñez, V., Pascual-Leone, A., Zhuang, P., and Hallett, M. (1998). Dynamic cortical involvement in implicit and explicit motor sequence learning. A PET study. Brain 121, 2159–2173.

Hunt, R.H., and Aslin, R.N. (2001). Statistical learning in a serial reaction time task: access to separable statistical cues by individual learners. J Exp Psychol Gen 130, 658–680.

Hutton, C., Bork, A., Josephs, O., Deichmann, R., Ashburner, J., and Turner, R. (2002). Image distortion correction in fMRI: a quantitative evaluation. Neuroimage 16, 217–240.

Jimenez, L. (2008). Taking patterns for chunks: is there any evidence of chunk learning in continuous serial reaction-time tasks? Psychol Res 72, 387–396.

Jimenez, L., Mendez, A., Pasquali, A., Abrahamse, E., and Verwey, W. (2011). Chunking by colors: assessing discrete learning in a continuous serial reaction-time task. Acta Psychol (Amst) 137, 318–329.

Jin, X., and Costa, R.M. (2010). Start/stop signals emerge in nigrostriatal circuits during sequence learning. Nature 466, 457–462.

Kass, R.E., and Raftery, A.E. (1995). Bayes Factors. Journal of the American Statistical Association 90, 773–795.

Kawashima, R., Matsumura, M., Sadato, N., Naito, E., Waki, A., Nakamura, S., Matsunami, K., Fukuda, H., and Yonekura, Y. (1998). Regional cerebral blood flow changes in human brain related to ipsilateral and contralateral complex hand movements–a PET study. European Journal of Neuroscience 10, 2254–2260.

Koch, I., and Hoffmann, J. (2000). Patterns, chunks, and hierarchies in serial reaction-time tasks. Psychol Res 63, 22–35.

Koechlin, E., and Jubault, T. (2006). Broca’s Area and the Hierarchical Organization of Human Behavior. Neuron 50, 963–974.

Kornysheva, K., and Diedrichsen, J. (2014). Human premotor areas parse sequences into their spatial and temporal features. Elife 3, e03043.

Kriegeskorte, N., Mur, M., and Bandettini, P. (2008). Representational similarity analysis - connecting the branches of systems neuroscience. Front Syst Neurosci 2, 4.

Kurata, K. (1991). Corticocortical inputs to the dorsal and ventral aspects of the premotor cortex of macaque monkeys. Neuroscience research 12, 263–280.

Lashley, K.S. (1951). The problem of serial order in behavior. In Cerebral mechanisms in behavior, pp. 112–136.

Margulies, D.S., Ghosh, S.S., Goulas, A., Falkiewicz, M., Huntenburg, J.M., Langs, G., Bezgin, G., Eickhoff, S.B., Castellanos, F.X., Petrides, M., et al. (2016). Situating the default-mode network along a principal gradient of macroscale cortical organization. Proceedings of the National Academy of Sciences 113, 12574–12579.

Margulies, D.S., Vincent, J.L., Kelly, C., Lohmann, G., Uddin, L.Q., Biswal, B.B., Villringer, A., Castellanos, F.X., Milham, M.P., and Petrides, M. (2009). Precuneus shares intrinsic functional architecture in humans and monkeys. Proc Natl Acad Sci U S A 106, 20069–20074.

Nambu, I., Hagura, N., Hirose, S., Wada, Y., Kawato, M., and Naito, E. (2015). Decoding sequential finger movements from preparatory activity in higher-order motor regions: a functional magnetic resonance imaging multi-voxel pattern analysis. European Journal of Neuroscience 42, 2851–2859.

Ng, A.Y., Jordan, M.I., and Weiss, Y. (2002). On spectral clustering: Analysis and an algorithm. Paper presented at: Advances in neural information processing systems.

Ohbayashi, M., Ohki, K., and Miyashita, Y. (2003). Conversion of working memory to motor sequence in the monkey premotor cortex. Science 301, 233–236.

Ohbayashi, M., Picard, N., and Strick, P.L. (2016). Inactivation of the Dorsal Premotor Area Disrupts Internally Generated, But Not Visually Guided, Sequential Movements. The Journal of neuroscience : the official journal of the Society for Neuroscience 36, 1971–1976.

Oosterhof, N.N., Wiestler, T., Downing, P.E., and Diedrichsen, J. (2011). A comparison of volume-based and surface-based multi-voxel pattern analysis. Neuroimage 56, 593–600.

Penhune, V.B., and Steele, C.J. (2012). Parallel contributions of cerebellar, striatal and M1 mechanisms to motor sequence learning. Behav Brain Res 226, 579–591.

Petit, L., Orssaud, C., Tzourio, N., Crivello, F., Berthoz, A., and Mazoyer, B. (1996). Functional anatomy of a prelearned sequence of horizontal saccades in humans. The Journal of neuroscience : the official journal of the Society for Neuroscience 16, 3714–3726.

Picard, N., and Strick, P.L. (1996). Motor areas of the medial wall: a review of their location and functional activation. Cereb Cortex 6, 342–353.

Ramkumar, P., Acuna, D.E., Berniker, M., Grafton, S.T., Turner, R.S., and Kording, K.P. (2016). Chunking as the result of an efficiency computation trade-off. Nat Commun 7, 12176.

Reber, A.S. (1967). Implicit learning of artificial grammars. Journal of Verbal Learning and Verbal Behavior 6, 855–863.

Rigoux, L., Stephan, K.E., Friston, K.J., and Daunizeau, J. (2014). Bayesian model selection for group studies - revisited. Neuroimage 84, 971–985.

Rosa, M.J., Bestmann, S., Harrison, L., and Penny, W. (2010). Bayesian model selection maps for group studies. Neuroimage 49, 217–224.

Rosenbaum, D.A., Kenny, S.B., and Derr, M.A. (1983). Hierarchical control of rapid movement sequences. J Exp Psychol Hum Percept Perform 9, 86–102.

Russo, A.A., Bittner, S.R., Perkins, S.M., Seely, J.S., London, B.M., Lara, A.H., Miri, A., Marshall, N.J., Kohn, A., and Jessell, T.M. (2018). Motor cortex embeds muscle-like commands in an untangled population response. Neuron 97, 953-966. e958.

Sadato, N., Campbell, G., Ibanez, V., Deiber, M., and Hallett, M. (1996). Complexity affects regional cerebral blood flow change during sequential finger movements. The Journal of neuroscience : the official journal of the Society for Neuroscience 16, 2691–2700.

Sakai, K., Hikosaka, O., Miyauchi, S., Takino, R., Sasaki, Y., and Pütz, B. (1998). Transition of brain activation from frontal to parietal areas in visuomotor sequence learning. Journal of Neuroscience 18, 1827–1840.

Sakai, K., Kitaguchi, K., and Hikosaka, O. (2003). Chunking during human visuomotor sequence learning. Exp Brain Res 152, 229–242.

Shen, S.M., W. J. (2018). Variable precision in visual perception. BioRxiv.

Shima, K., Isoda, M., Mushiake, H., and Tanji, J. (2007). Categorization of behavioural sequences in the prefrontal cortex. Nature 445, 315–318.

Stadler, M.A. (1992). Statistical Structure and implicit serial learning. Journal of Experimental Psychology: Learning, Memory and Cognition 18, 318–327.

Stephan, K.E., Penny, W.D., Daunizeau, J., Moran, R.J., and Friston, K.J. (2009). Bayesian model selection for group studies. Neuroimage 46, 1004–1017.

Tanji, J., and Shima, K. (1994). Role for supplementary motor area cells in planning several movements ahead. Nature 371, 413–416.

Tomassini, V., Jbabdi, S., Klein, J.C., Behrens, T.E.J., Pozzilli, C., Matthews, P.M., Rushworth, M.F.S., and Johansen-Berg, H. (2007). Diffusion-Weighted Imaging Tractography-Based Parcellation of the Human Lateral Premotor Cortex Identifies Dorsal and Ventral Subregions with Anatomical and Functional Specializations. The Journal of Neuroscience 27, 10259–10269.

Verwey, W.B., and Abrahamse, E.L. (2012). Distinct modes of executing movement sequences: reacting, associating, and chunking. Acta Psychol (Amst) 140, 274–282.

Verwey, W.B., and Dronkert, Y. (1996). Practicing a structured continuous key-pressing task: Motor chunking or rhythm consolidation? Journal of motor behavior 28, 71–79.

Von Luxburg, U. (2007). A tutorial on spectral clustering. Statistics and computing 17, 395–416.

Walther, A., Nili, H., Ejaz, N., Alink, A., Kriegeskorte, N., and Diedrichsen, J. (2016). Reliability of dissimilarity measures for multi-voxel pattern analysis. Neuroimage 137, 188–200.

Wiestler, T., and Diedrichsen, J. (2013). Skill learning strengthens cortical representations of motor sequences. Elife 2, e00801.

Wiestler, T., Waters-Metenier, S., and Diedrichsen, J. (2014). Effector-independent motor sequence representations exist in extrinsic and intrinsic reference frames. The Journal of neuroscience : the official journal of the Society for Neuroscience 34, 5054–5064.

Wymbs, N.F., Bassett, D.S., Mucha, P.J., Porter, M.A., and Grafton, S.T. (2012). Differential recruitment of the sensorimotor putamen and frontoparietal cortex during motor chunking in humans. Neuron 74, 936–946.

Yamashita, Y., and Tani, J. (2008). Emergence of functional hierarchy in a multiple timescale neural network model: a humanoid robot experiment. PLoS Comput Biol 4, e1000220.

Yeo, B.T.T., Krienen, F.M., Sepulcre, J., Sabuncu, M.R., Lashkari, D., Hollinshead, M., Roffman, J.L., Smoller, J.W., Zöllei, L., Polimeni, J.R., et al. (2011). The organization of the human cerebral cortex estimated by intrinsic functional connectivity. Journal of Neurophysiology 106, 1125–1165.

Yokoi, A., Arbuckle, S.A., and Diedrichsen, J. (2018). The Role of Human Primary Motor Cortex in the Production of Skilled Finger Sequences. The Journal of neuroscience : the official journal of the Society for Neuroscience 38, 1430–1442.

Yokoi, A., Bai, W., and Diedrichsen, J. (2017). Restricted transfer of learning between unimanual and bimanual finger sequences. J Neurophysiol 117, 1043–1051.

